# A Protein Kinase A-Regulated Network Encodes Short- and Long-Lived Cellular Memory

**DOI:** 10.1101/656009

**Authors:** Yanfei Jiang, Zohreh AkhavanAghdam, Yutian Li, Brian M. Zid, Nan Hao

## Abstract

Cells can store memories of prior experiences to modulate their responses to subsequent stresses, as seen for the protein kinase A (PKA)-mediated general stress response in yeast, which is required for resistance against future stressful conditions. Using microfluidics and time-lapse microscopy, we quantitatively analyzed how cellular memory of stress adaptation is encoded in single yeast cells. We found that cellular memory is biphasic. Short-lived memory was mediated by trehalose synthase and trehalose metabolism. Long-lived memory was mediated by PKA-regulated stress-responsive transcription factors and cytoplasmic messenger ribonucleoprotein (mRNP) granules. Strikingly, short- and long-lived memory could be selectively induced by different priming input dynamics. Computational modeling revealed how the PKA-mediated regulatory network could encode previous stimuli into memories with distinct dynamics. This biphasic memory-encoding scheme, analogous to innate and adaptive immune responses in mammals, might represent a general strategy to prepare for future challenges in rapidly changing environments.

## Introduction

Cells survive rapidly changing environments through adaptation mediated by sophisticated signaling and gene regulatory networks. How these networks operate dynamically to process complex extracellular signals and elicit appropriate responses remains a challenging question (Behar and Hoffmann, 2010; Levine et al., 2013; Purvis and Lahav, 2013). Recent advances in microfluidics and single-cell imaging technologies allow us to track the responses of individual living cells in a precisely controlled changing environment, providing a unique opportunity to elucidate the underlying principles for dynamic signal processing in cells (Bennett and Hasty, 2009). In this study, we exploited these cutting-edge technologies to systematically probe the regulatory network that enables cells to encode memory of prior environmental cues in order to modulate their adaptive responses to future challenges.

History-dependent cellular behaviors have been found in many organisms (Durrant and Dong, 2004; Hilker et al., 2016; Lou and Yousef, 1997; Matsumoto et al., 2007; Netea et al., 2015; Schenk et al., 2000; Scholz et al., 2005). For instance, it has been long established in the field of plant physiology that plant cells, once primed by mild stresses or chemical compounds, could obtain enhanced resistance to various future diseases and abiotic stresses (Prime et al., 2006; Savvides et al., 2016). In another oft-cited example, pre-treatment of human macrophages with interferon-γ significantly boosts subsequent lipopolysaccharide (LPS) responses against various pathogens and in tumor cell killing (Gifford and Lohmann-Matthes, 1987; Hayes et al., 1995). In this study, we referred these history-dependent responses in single cells as “cellular memory”, which is of course fundamentally different from the neuronal memory in animals.

In the yeast *Saccharomyces cerevisiae*, a given stress can activate its specific response pathway as well as a common signaling pathway shared by a wide range of different stresses, called the general stress response (GSR) pathway (Gasch et al., 2000). This pathway is primarily mediated by protein kinase A (PKA). In response to stresses, PKA is rapidly inhibited, leading to activation of downstream transcription factors (TFs), such as Msn2 and Msn4, and the induction of hundreds of stress responsive genes (Gorner et al., 1998; Hao and O’Shea, 2012; Jiang et al., 2017; Martinez-Pastor et al., 1996). Previous studies have demonstrated that GSR is not required for survival against immediate stress threats, but instead, is required for resistance against future stressful conditions (Berry and Gasch, 2008; Berry et al., 2011; Guan et al., 2012). However, the mechanisms that mediate memory encoding of environmental changes remain unclear.

In this study, we used GSR as a model to quantitatively analyze how PKA-dependent regulatory processes operate dynamically to encode the memory of environmental changes. We combined microfluidics with time-lapse microscopy to precisely control the dynamics of priming inputs and to quantify the memory effect on stress adaptation in single cells. We found that cellular memory shows two phases, a fast-decaying phase mediated by trehalose metabolism and a long-lasting phase mediated by stress-activated TFs and messenger ribonucleoprotein (mRNP) granules. Moreover, the memory dynamics can be modulated by priming inputs. Whereas a high amplitude transient input specifically induces the fast-decaying phase of memory, a prolonged input is needed to elicit the long-lasting memory effect. We further developed a computational model based on the molecular processes identified experimentally. Our model quantitatively revealed the regulatory scheme that encodes the information of previous environmental inputs into distinct memory dynamics, implying a general strategy to optimize resource allocation and prepare for future challenges under rapidly changing environments.

## Results

### PKA encodes biphasic cellular memory

Probing the effect of cellular memory has long been challenging, in part due to the difficulty in generating well-controlled sequential environmental changes in cell cultures. Recent advances in microfluidic technologies allow the precise control of extracellular conditions and tracking of the responses of single cells over extended periods (Hansen et al., 2015; Li et al., 2017a), providing a powerful tool for analyzing the memory behaviors in stress responses. In this study, we modified the channel design of an existing microfluidic device (Hao and O’Shea, 2012) to include separate control of three media inlets, one for the priming input, one for the normal growth medium, and one for the stress treatment (Fig. 1A). To increase the experimental throughput, we further aligned four individual channels into a single device to enable simultaneous running of multiple experiments, each with its own cell population and stimulus condition. Using this device, we first exposed the cells to a pulse of priming input followed by a “break time” with the normal growth medium. We then treated these primed cells with a sustained environmental stress and evaluated their adaptation responses. The device was mounted on a time-lapse microscope to track the responses of a large number of single cells throughout the entire experiment.

**Figure 1.**
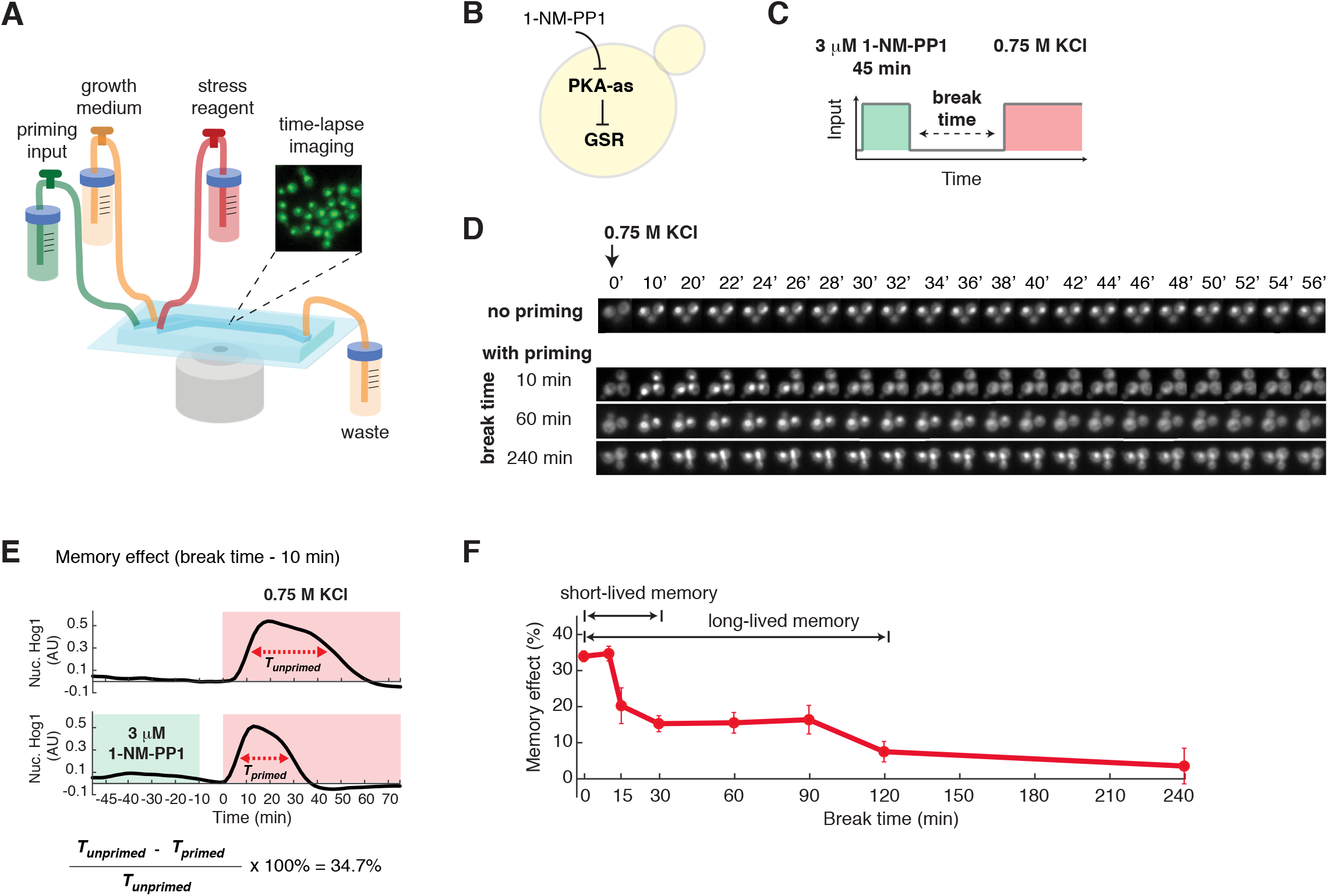
PKA encodes biphasic cellular memory. (A) A diagram of the microfluidic system used in priming experiments. The microfluidic device contains three inlets for priming input, growth medium, and stress reagent, respectively. Yeast cells were immobilized in the culturing chamber for time-lapse microscopic imaging. The medium flow from the inlets to the waste is driven by gravity. The three inlets can be opened/closed to allow medium switching during the experiment. (B) A diagram describes the analog sensitive PKA strain used in this study. (C) A schematic of the experimental design. Within the microfluidic device, cells were first exposed to a pulse of 1-NM-PP1 priming input, followed by a break time of normal growth medium. These primed cells were then exposed to sustained 0.75 M KCl treatment. (D) Representative time-lapse images of Hog1 nuclear translocation in the priming experiments with different break times. Left panel - schematic illustration of the priming experiments. (E) Time traces of Hog1 translocation in response to 0.75 M KCl (red shaded) without (top) or with (bottom) 45 min priming with 3 µM 1-NM-PP1 (green shaded) followed by 10 min break time. The duration of nuclear localization was quantified using the full width at half maximum (FWHM) in single cells (Tprimed and Tunprimed). The memory effect has been calculated using Tprimed and Tunprimed. (F) Biphasic memory dynamics in response to the high-amplitude prolonged priming input (panel C). The plot shows the relationship of memory effect versus break time. Data points are averages of at least three independent experiments. Error bars – standard error of the mean (SEM). The Hog1 time trace for each break time is shown in Figure S1.

For the priming input, we used a chemical genetics strategy in which we introduced analog-sensitive mutations into the PKA isoforms so that they remain fully functional but can be specifically inhibited by the small molecular inhibitor, 1-NM-PP1 (Bishop et al., 2000). We have previously employed this strategy to control PKA activity, mimicking an upstream signaling event that specifically activates GSR, but not other stress specific pathways (AkhavanAghdam et al., 2016; Hansen and O’Shea, 2013; Hao et al., 2013; Hao and O’Shea, 2012) (Fig. 1B). Moreover, combined with time-lapse microscopy and microfluidics, it enables us to generate precisely controlled temporal patterns of PKA inhibition as priming inputs and evaluate their effects on cells’ adaptation to the subsequent environmental stress.

For the subsequent stress treatment, we chose hyperosmotic stress (0.75M KCl) because the stress adaptation process in individual cells can be reliably quantified using a specific reporter, the stress-activated protein kinase Hog1, which we tagged with YFP. In response to osmotic stress, Hog1 rapidly translocates to the nucleus to induce an increase in intracellular osmolyte; once the osmolyte balance is restored and the cell recovers from the stress, Hog1 exits the nucleus (Brewster and Gustin, 2014). There is a strong correlation between the timing of Hog1 nuclear export and the restoration of cell volume (reflecting the turgor pressure recovery and cellular adaptation) (Muzzey et al., 2009). Thus, the duration of Hog1 nuclear localization serves as a proxy for the time needed for the cell to recover from a stress treatment: a shorter duration represents a faster adaptation while a longer duration represents a slower adaptation.

Using Hog1 nuclear localization as a reporter, we observed that a 45-min priming input with 3 µM PKA inhibitor (Fig. 1C), which causes full PKA inhibition (Hao et al., 2013; Hao and O’Shea, 2012), dramatically shortened the time needed to recover from hyperosmotic stress. Furthermore, the effect of this priming input decays with increasing break times (Fig. 1D). To quantify the effect of priming input and the dynamics of its decay, we measured and compared the durations of Hog1 nuclear translocation with and without the priming input (*T*_*primed*_, *T*_*unprimed*_) for each break time. For instance, the priming input with a 10-min break time decreased the adaptation time by 34.7% from 40 min to 26min (Fig.1E).We defined this percentage decrease in recovery 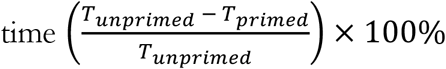 as a quantitative measure of the “memory effect” and use it throughout our report.

We then plotted the memory effect as a function of break time. Notably, we observed that the decay of memory effect is biphasic: About half of the memory effect was lost rapidly within 30 minutes, whereas the remaining memory effect plateaued until 90 minutes and then declined slowly (Fig. 1F; Fig. S1). These biphasic memory dynamics were in contrast with the scenario where the memory is primarily mediated by the expression of stress-resistance genes (Berry and Gasch, 2008; Guan et al., 2012), which would follow an exponential decay trajectory as gene products are degraded during the break time. We termed the fast-decaying component “short-lived” memory and the longer-lasting component “long-lived” memory. To test whether the memory effect shows two phases under natural stress conditions, we used 0.5M KCl as the priming input and observed similar biphasic memory dynamics (Fig. S2), confirming that the memory pattern we found is not specific to chemical inhibition of PKA.

### Short- and long-lived memory can be selectively induced by different priming input dynamics

We next considered how changing the dynamics of priming inputs impacts cellular memory. To examine the effect of input duration on cellular memory, we exposed cells to a 15-min priming input with 3 µM 1-NM-PP1 (Fig. 2A). In contrast to the response upon a 45-min priming input with the same amplitude, cells only exhibited the fast-decaying component of memory effect and the long-lasting phase is completely abolished (Fig. 2B, compare the blue versus dark pink curves). When cycloheximide was added to inhibit protein synthesis during the priming experiment, this short-lived memory effect was not affected (Fig. 2C), suggesting that short-lived memory does not depend on gene expression, but may instead be mediated by metabolites or post-translational protein modifications.

**Figure 2.**
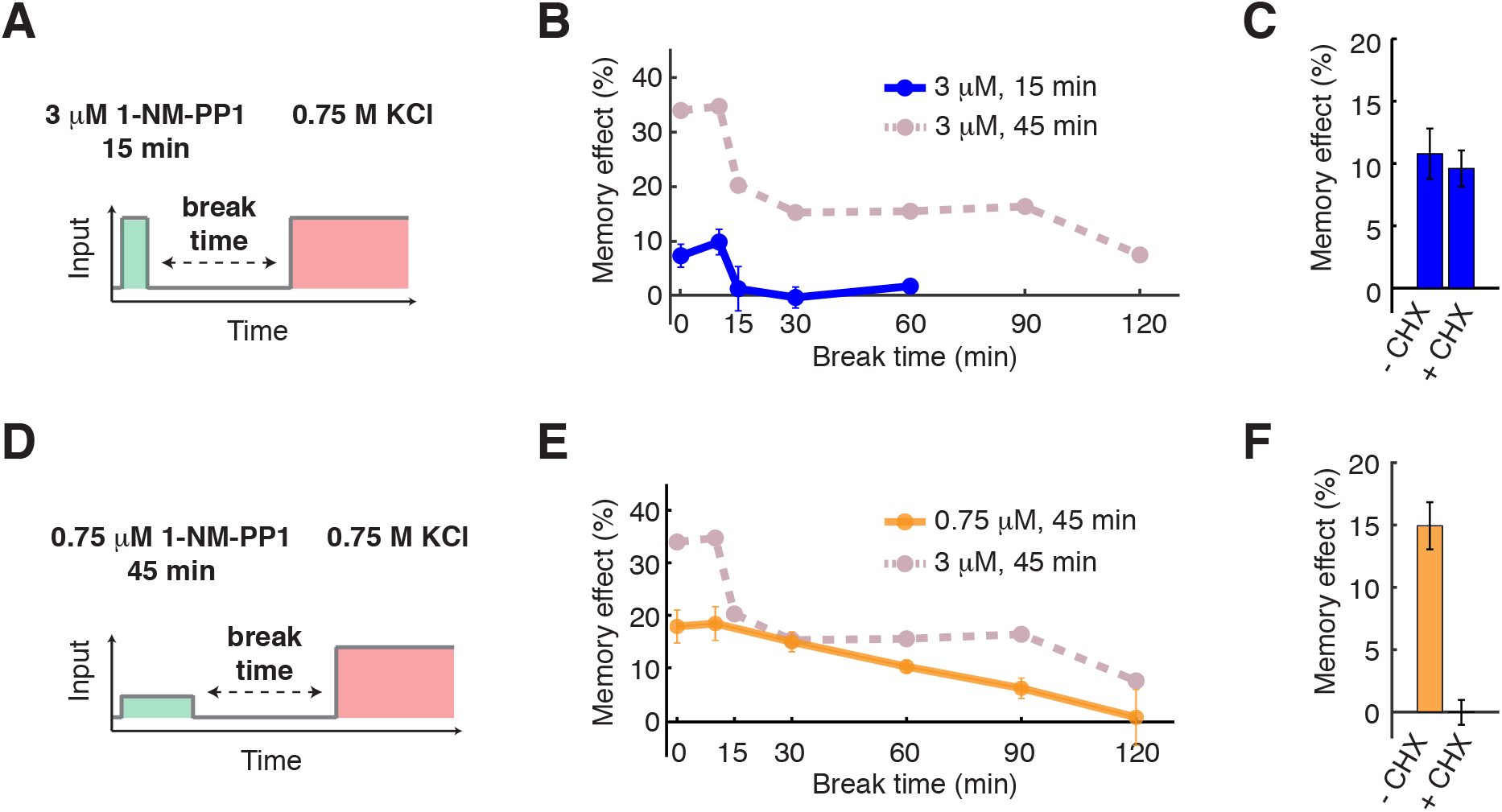
Short- and long-lived memory can be selectively induced by different priming input dynamics. (A) A schematic of the priming experiment with the high-amplitude transient priming input (15 min, 3 µM 1-NM-PP1). (B) The dynamics of memory effect in response to the high-amplitude transient priming input. Dashed line in dark pink represents the memory dynamics from Fig. 1F and is included in the plot for comparison. (C) The dependence of short-lived memory effect on translation. The bar graph shows the memory effects in response to 15 min, 3 *µ*M 1-NM-PP1 priming input with a 10 min break time, with or without 10 *µ*g/mL cycloheximide treatment (CHX). (D) A schematic of the priming experiment with the low-amplitude prolonged priming input (45 min, 0.75 *µ*M 1-NM-PP1). (E) The dynamics of memory effect in response to the low-amplitude prolonged priming input. Dashed line in dark pink represents the memory dynamics from Fig. 1F and is included in the plot for comparison. (F) The dependence of long-lived memory effect on translation. The bar graph shows the memory effects in response to 45 min, 0.75 *µ*M 1-NM-PP1 priming input with a 30 min break time, with or without the CHX treatment. Data points are averages of at least three independent experiments. Error bars – SEM.

To further evaluate the effect of input amplitude on cellular memory, we treated cells with a 45-min priming input with lower amplitude (0.75 µM 1-NM-PP1), which causes partial inhibition of PKA (Hao et al., 2013; Hao and O’Shea, 2012)(Fig. 2D). Under this condition, the fast-decaying memory component and the plateau phase (0 – 90 minutes) of long-lived memory were both abolished (Fig. 2E, compare the orange versus dark pink curves). Instead, cells displayed a slow exponential decay trajectory, as would be expected if the memory effect is primarily mediated by gene expression. In accordance, CHX abolished this memory effect (Fig. 2F), confirming that it requires gene expression.

Taken together, these results demonstrated that the short- and long-lived components of cellular memory could be dissected by modulating the amplitude and duration of priming inputs and might be mediated by different molecular processes. Short-lived memory might be mediated through a fast translation-independent process, whereas an input duration-dependent slow gene expression process mediates the long-lasting memory, which could be further stabilized by another input amplitude-dependent mechanism. The three types of priming inputs used here are referred to as: “high-amplitude prolonged” (3 µM 1-NM-PP1, 45 min); “high-amplitude transient” (3 µM, 15 min); and “low-amplitude prolonged” (0.75 µM, 45 min).

### Short-lived memory is mediated by trehalose synthesis and metabolism

To determine the molecular process that mediates short-lived memory, we systematically deleted 5 known or putative PKA phosphorylation targets (Gph1, Ctt1, Nth1, Gcy1, and Tps1) involved in metabolic or stress-response pathways (Hwang et al., 1989; Schepers et al., 2012; Trevisol et al., 2014) and examined their roles in mediating memory effect. Among these PKA targets, we identified the trehalose synthase Tps1 (De Virgilio et al., 1993; Reinders et al., 1997), deletion of which abolished short-lived memory (Fig. 3A). Trehalose is a simple carbohydrate produced in many organisms that acts as membrane protectant and protein stabilizer to enhance cell survival under stressful conditions (Hounsa et al., 1998). Note that the *tps1*Δ*hxk2*Δstrain was used because *tps1*Δis unable to grow on glucose, but this growth is restored in a *tps1*Δ*hxk2*Δdouble mutant (Hohmann et al., 1993). The deletion of *HXK2* alone does not affect the memory effect (Fig. 3A). We further confirmed that the intracellular trehalose level is rapidly increased in response to the PKA inhibition input and that this increase is lost when Tps1 is deleted (Fig. 3B). Since trehalose has been shown to play an important role in cellular protection against osmotic stress (Hounsa et al., 1998), the increased level of trehalose by the priming input could temporarily enhance the acquired resistance to the subsequent osmotic stress, accounting for the observed short-lived memory.

**Figure 3.**
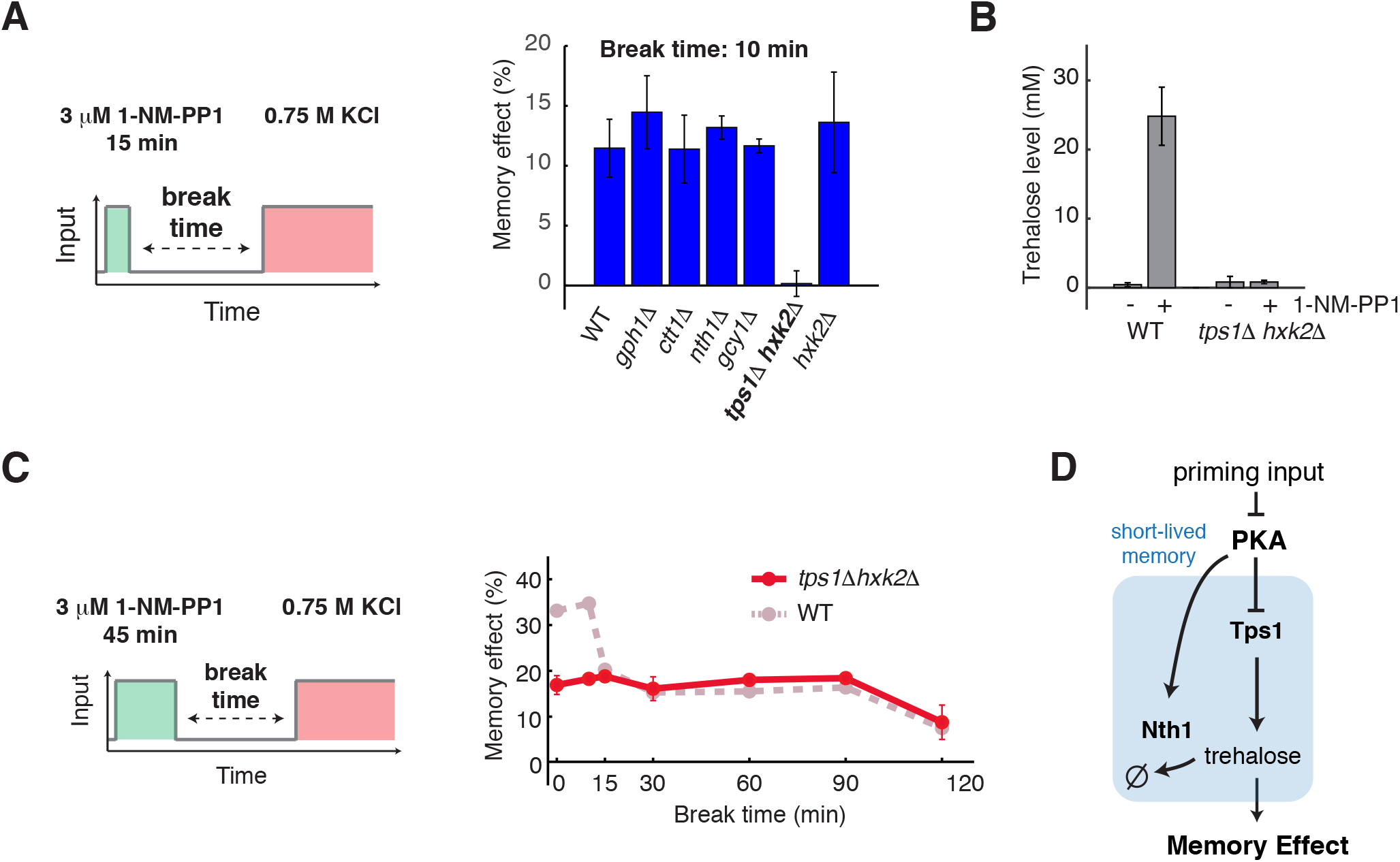
Short-lived memory is mediated by trehalose synthesis and metabolism. (A) Identification of Tps1 in mediating short-term memory. The bar graph shows the memory effects in response to the high-amplitude transient priming input (15 min, 3 *µ*M 1-NM-PP1) with a 10 min break time in WT and mutant strains. (B) Intracellular trehalose level increases in response to PKA inhibition. Bar graph shows trehalose levels in WT and *tps1*Δ*hxk2Δ*, with and without PKA inhibition. Trehalose levels were measured after a 20-min treatment of 3 µM 1-NM-PP1. (C) Memory dynamics in *tps1*Δ*hxk2*Δin response to the high-amplitude prolonged priming input (45 min, 3 µM 1-NM-PP1). Left panel, the schematic illustrating the treatment procedure of the priming experiment. Right panel, memory dynamics in *tps1*Δ*hxk2Δ*. Dashed lines in dark pink represent the memory dynamics in WT (from Fig. 1F) and are included in the plots for comparison. Data points are averages of at least three independent experiments. Error bars – SEM. (D) A diagram illustrates the network motif that gives rise to short-lived memory.

To validate the central role of trehalose in short-lived memory, we manipulated its degradation. Nth1 is a trehalose degradation enzyme, the activity of which is also regulated by PKA-dependent phosphorylation (Schepers et al., 2012; Souza et al., 2002). We observed that, whereas short-term memory was abolished in the absence of Tps1, it became prolonged when *NTH1* is deleted (Fig. S3, compare the blue versus grey blue curves), suggesting that the activation of Nth1 accounts for the fast decay rate of the trehalose level. In other words, the duration of short-lived memory is encoded by the activity of Nth1.

To test whether Tps1 and trehalose also contribute to long-lived memory, we examined the memory dynamics in response to the high-amplitude prolonged input (3 µM 1-NM-PP1, 45 min). We found that the absence of Tps1 completely abolished the fast-decaying component of memory, while long-lived memory remained unchanged (Fig. 3C, compare the red versus dark pink curves), indicating that trehalose synthesis and metabolism contribute exclusively to short-lived memory. In summary, these results showed that short-lived memory is mediated by PKA-dependent regulation of Tps1 and Nth1, which control trehalose synthesis and metabolism (Fig. 3D).

### Long-lived memory is mediated by stress-activated TFs and mRNP granules

We next investigated the processes underlying long-lived memory, which is gene expression-dependent, as suggested above in Fig. 2F. To determine the TFs that induce this response, we systematically deleted 6 PKA-regulated stress-responsive TFs (Gis1, Sko1, Hot1, Yap1, and Msn2/4) (Charizanis et al., 1999; Gorner et al., 1998; Hinnebusch and Natarajan, 2002; Pascual-Ahuir et al., 2001; Proft et al., 2001; Roosen et al., 2005) and examined their roles in mediating memory effect. We found that, whereas the deletion of Msn2/4 completely abolished long-lived memory in response to the 45 minutes, 0.75 µM 1-NM-PP1 priming input (Fig. S4A), the mutant showed only a partial loss of memory in response to the 3 µM 1-NM-PP1 priming input (Fig. S4B), suggesting that another TF might play a compensatory role under this condition. As the deletion of Yap1 also dramatically diminished long-lived memory in response to the 0.75 µM 1-NM-PP1 priming input (Fig. S4A), we generated the *msn2*Δ*msn4*Δ*yap1*Δtriple mutant and observed a complete loss of long-lived memory in the triple mutant (Fig. S4B). These results identified Msn2/4 and Yap1 as primary TFs that mediate the transcriptional response generating long-lasting memory. Moreover, we observed that the short- and long-lived memories are both abolished in the triple mutant (Fig. 4A and B). This loss of short-lived memory in the triple mutant is consistent with previous studies (Hansen and O’Shea, 2013; Norbeck and Blomberg, 2000), which showed that the expression of *TPS1*, required for short-lived memory, is completely dependent on the TFs deleted in the triple mutant.

**Figure 4.**
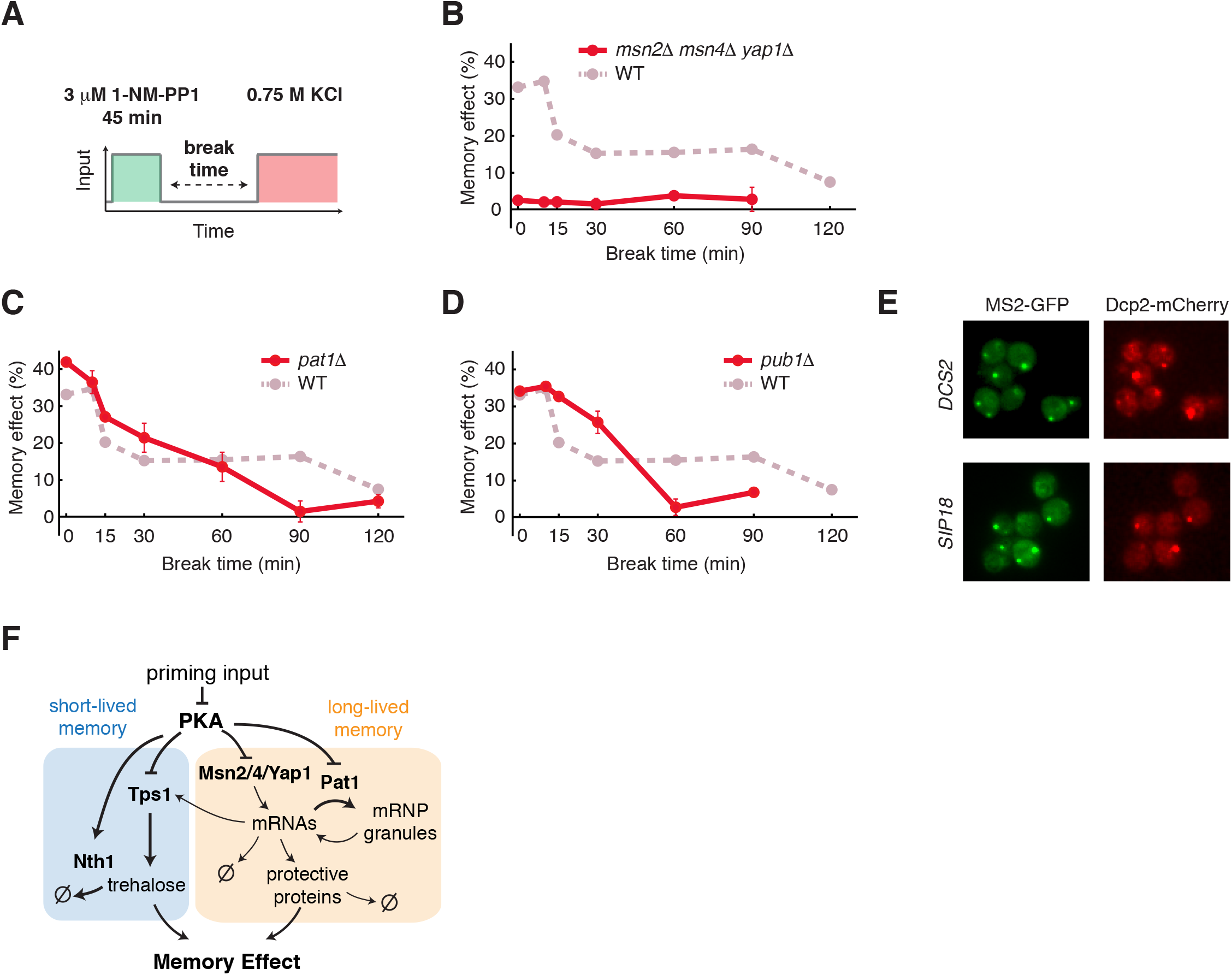
Long-lived memory is mediated by stress-activated TFs and mRNP granules. (A) A schematic of the priming experiment with the high-amplitude prolonged priming input (45 min, 3 µM 1-NM-PP1). Memory dynamics in (B) *msn2*Δ*msn4*Δ*yap1Δ*, (C) *pat1Δ*, and (D) *pub1Δ*in response to the high-amplitude prolonged priming input. Data points are averages of at least three independent experiments. Error bars – SEM. Dashed line in dark pink represents the memory dynamics in WT (from Fig. 1F) and is included in the plot for comparison. (E) Co-localization of *DCS2* or *SIP18* mRNAs with PBs (Dcp2-mCherry) in response to 3 µM 1-NM-PP1. Foci formation was observed within 20 min of the input treatment. Representative images were acquired at 170 min. (F) A diagram illustrates the PKA-regulated network that mediates short-lived and long-lived memory.

We then investigated the mechanism that stabilizes long-lived memory, underlying the plateau phase of memory that is maintained up to 90 minutes after the removal of priming inputs (Fig. 1F). Previous studies have revealed that, in response to stress or PKA inhibition, cells accumulate a number of stress-responsive gene mRNAs in cytoplasmic mRNP granules, called processing-bodies (PBs) and stress granules (SGs), which regulate mRNA translation, decay, and storage (Decker and Parker, 2012; Ramachandran et al., 2011; Wang et al., 2018; Zid and O’Shea, 2014). PKA regulates the formation of these mRNP granules by phosphorylating a PB scaffolding protein, Pat1 (Ramachandran et al., 2011). To test whether mRNP granules contribute to the memory effect, we deleted Pat1, which is essential for the formation of PBs (Buchan et al., 2008; Ramachandran et al., 2011). We observed that the plateau phase of long-lived memory was abolished in the *pat1*Δstrain and the memory effect exhibited a continuous decay (Fig. 4C, compare the red versus dark pink curves), indicating that PBs are required for maintaining the plateau phase. PBs and SGs are discrete but functionally interacting compartments. Some mRNAs in PBs can be stored in a translationally silenced state during stress and then return to translation via SGs upon stress removal (Buchan et al., 2008; Decker and Parker, 2012). To determine the role of SGs in memory, we examined memory dynamics in the absence of the SG component Pub1 (Buchan et al., 2008). We observed that, similar to *pat1Δ, pub1Δ*cells no longer exhibited the plateau phase of memory (Fig. 4D, compare the red versus dark pink curves). We note that *pub1Δ*cells have similar growth rate to that of WT throughout the experiments, indicating that the plateau phase observed in WT is not related to cell growth rate. These results suggested that the plateau phase of long-lived memory is dependent on the PBs/SGs-mediated mRNA storage pathway. Notably, *pat1*Δand *pub1Δ*showed higher memory levels than those of WT for shorter break times (Fig. 4C and D, 15 - 30 mins), in agreement with the role of PBs/SGs in mRNA translational silencing in addition to storage.

Furthermore, to confirm the localization of the mRNAs of stress responsive genes to mRNP granules, we used the MS2 coat protein (MS2-CP) fused to GFP (Bertrand et al., 1998; Haim-Vilmovsky and Gerst, 2009) to visualize the mRNAs of two well-characterized PKA-regulated stress responsive genes, *DCS2* and *SIP18* (AkhavanAghdam et al., 2016; Hansen and O’Shea, 2013) in living cells. We observed that, in response to the 3 µM 1-NM-PP1 input, the mRNAs of *DCS2* and *SIP18* formed foci that co-localized with PBs as indicated by the PB marker Dcp2-mCherry (Brengues et al., 2005) as well as some distinct granules (Fig. 4E). We have previously observed a similar localization pattern for Msn2/4 targets, such as *GLC3* and *HXK1*, during glucose starvation (Zid and O’Shea, 2014). This localization pattern coincided with poor protein production from these mRNAs during stress, suggesting that these stress-induced mRNAs are translationally silenced and stored in mRNP granules to confer long-lasting cellular memory.

Taken together, these findings uncovered that two processes, gene transcription and mRNA storage by mRNP granules, operate together to generate long-lived memory (Fig. 4F).

### Computational modeling suggested the network-mediated encoding of memory dynamics

From the experimental analysis described above, we delineated the regulatory processes that mediate cellular memory. To quantitatively understand the dynamic encoding of memory, we constructed a computational model. In the model, the network is composed of two memory-encoding motifs, one for short-lived memory and the other for long-lived memory (Fig. 4F). The short-lived memory motif comprises a fast process, in which PKA regulates the activities of Tps1 and Nth1 by phosphorylation (Schepers et al., 2012; Trevisol et al., 2014). In response to PKA inhibition, Tps1 is activated, boosting trehalose production; at the same time, Nth1 is inhibited, slowing down trehalose degradation. As a result, the trehalose level increases rapidly. Subsequently, when the PKA inhibition input is removed, Tps1 is inhibited while Nth1 is activated, leading to a rapid decline in trehalose levels during the break time. This feedforward loop enables quick tuning of trehalose levels, accounting for the fast-changing component of memory effect. The long-lived memory motif consists of two processes that function together to regulate gene expression dynamics, a transcriptional response mediated by TFs Yap1 and Msn2/4 (Smith et al., 1998) and a mRNA storage process mediated by Pat1 (Decker and Parker, 2012). A detailed description of the model is included in the Supplemental Materials.

With computationally obtained best-fit parameters, our model reproduced all the experimental data (Fig. S5). In particular, in our model, the high-amplitude transient input (Fig. 5A, left panel) specifically induces the trehalose production process (Fig. 5A, middle panel), generating the fast-decaying memory (Fig. 5A, right panel, compare with data in Fig. 2B). The low-amplitude prolonged input (Fig. 5B, left panel) only induce the transcriptional response but not the mRNP granule formation, because the input is too weak, (Fig. 5B, middle panel), resulting in a slow exponential decay of the memory effect after input removal (Fig. 5C, right panel, compare with data in Fig. 2E). The high-amplitude prolonged input (Fig. 5C, left panel), however, leads to co-activation of the fast trehalose production and the slow transcriptional response with mRNA storage by mRNP granules (Fig. 5C, middle panel). Once the transcriptional response is initiated, a portion of the newly synthesized stress resistance gene mRNAs is stored in mRNP granules via PKA regulation of Pat1. Upon input removal, these mRNAs are slowly released and translated, resulting in a long-lasting (up to ∼90 minutes) memory plateau (Fig. 5C, right panel, compare with data in Fig. 1F). Consistent with the model, we observed that only the high-amplitude prolonged input strongly induced the formation of mRNP granules, as indicated by *DCS2* mRNA foci, whereas the high-amplitude transient input or low-amplitude prolonged input could not (Fig. 5, right panel insets). These live-cell mRNA results support our model in which the formation and function of mRNP granules depend on both the input amplitude and duration. In summary, our model suggested that the three PKA-regulated processes, trehalose metabolism, gene transcription and mRNP granule formation, operate and coordinate in a temporal order to enable the biphasic memory dynamics observed experimentally.

**Figure 5.**
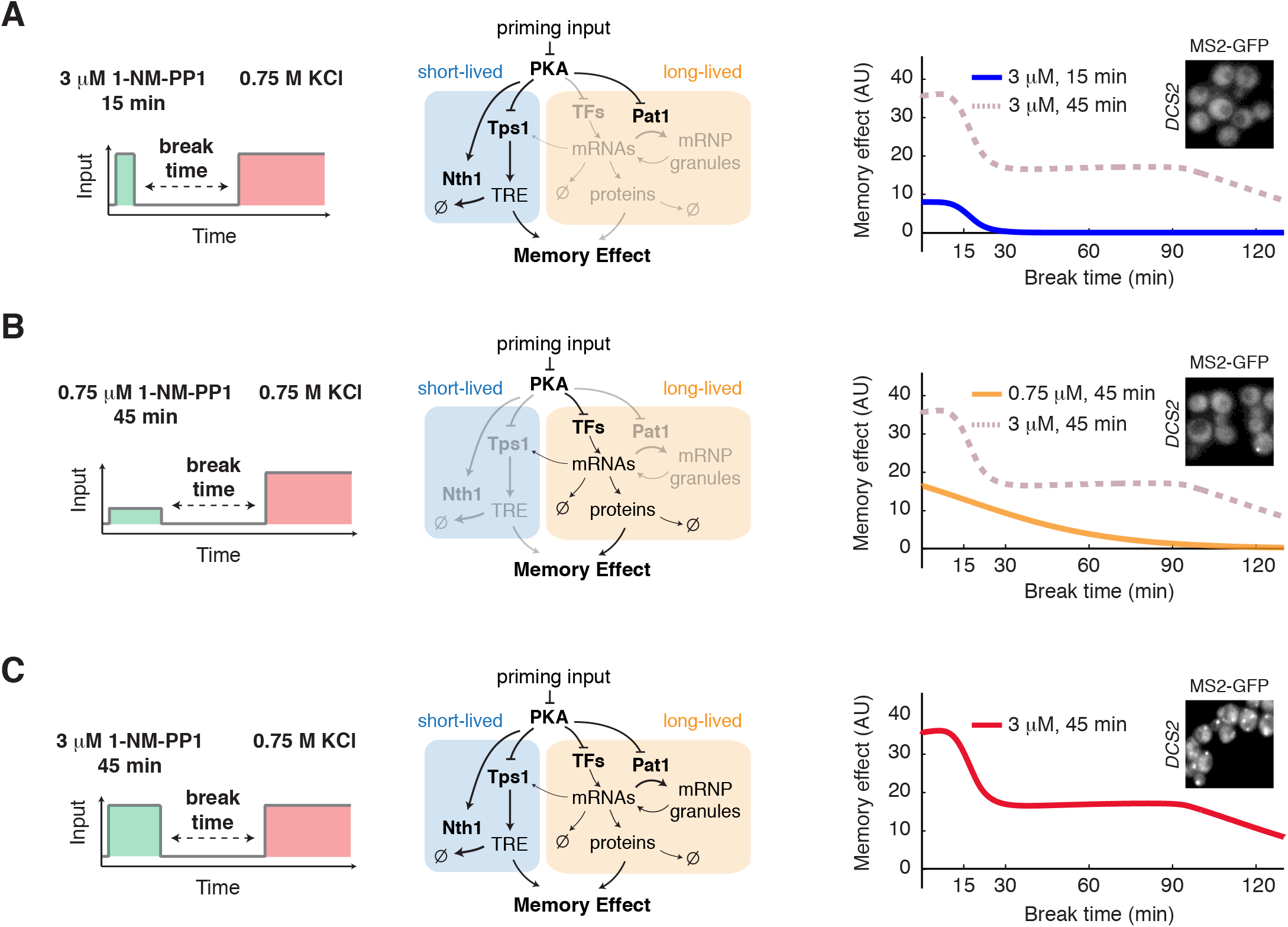
Computational modeling reveals the network-mediated encoding of cellular memory. (A) Model simulation of memory dynamics in response to the high-amplitude transient priming input (15 min, 3 *µ*M 1-NM-PP1). Left panel, the schematic illustrates the treatment procedure. Middle panel, the diagram highlights the part of the network that is activated by the input (black – activated; light gray – not activated). Right panel, the plot shows the simulated dynamics of memory effect. Dashed line in dark pink represents the simulated dynamics in response to the high-amplitude prolonged priming input (45 min, 3 *µ*M 1-NM-PP1) and is included in the plot for comparison. (B) Model simulation of memory dynamics in response to the low-amplitude prolonged priming input (45 min, 0.75 *µ*M 1-NM-PP1). (C) Model simulation of memory dynamics in response to the high-amplitude prolonged priming input (45 min, 3 *µ*M 1-NM-PP1). Insets, the granule formation of DCS2 mRNA in response to indicated inputs. Representative images were acquired at the end of input treatments.

Moreover, our model could nicely reproduce the memory dynamics observed in the mutants of key regulatory factors. In the absence of Tps1, the priming input cannot induce the production of trehalose (Fig. S6A, middle panel), resulting in a loss of the fast decaying component of memory, while the long-lived memory remains intact (Fig. S6A, right panel, compare with data in Fig. 3C). When the TFs Msn2/4 and Yap1 are deleted, the priming input can no longer induce the expression of stress resistance genes, resulting in a loss of long-lived memory. Meanwhile, the absence of these TFs leads to the loss of Tps1 expression that is required for the short-lived memory (Fig. S6B, middle panel). As a result, both short- and long-lived memories are completely abolished (Fig. S6B, right panel, compare with data in Fig. 4A). When Pat1 is deleted, the newly synthesized mRNAs, induced by the priming input, can no longer be stored in mRNP granules (Fig. S6C, middle panel), but instead they undergo immediate translation, resulting in a higher initial level of stress resistance than that of WT (during break time 0 – 30 min). This stress resistance, however, decays more quickly, as the gene products are being directly and continuously degraded, and cells can no longer maintain a plateau of long-lived memory (Fig. S6C, right panel, compare with data in Fig. 4C).

### Model prediction and experimental validation

To further test our model, we used it to predict the memory dynamics in response to a combined pattern of priming input. Neither low-amplitude prolonged input nor high-amplitude transient input can induce the plateau phase of memory (Fig. 2B and E), yet our model predicted that these two inputs, when applied sequentially (Fig. 6A), should be capable of generating a memory plateau phase that is Pat1 dependent (Fig. 6B, “Prediction”). In this scenario, the low-amplitude prolonged input would first produce a high level of mRNAs, and then the subsequent high-amplitude transient input would induce mRNP granules to store newly synthesized mRNAs, enabling the plateau phase. This prediction is intriguing because it illustrates that the memory effect to the combined input is not simply the sum of the effects to the two single inputs (Fig. 6B, “Prediction”, compare the purple versus light pink curves) and this is attributed to the mRNP-dependent storage mechanism. Therefore, since the memory-encoding processes are largely independent, when the mRNA storage process is removed in the *pat1Δ*mutant, the memory effects would become additive (Fig. 6C, “Prediction”). We experimentally tested these predictions, and, in agreement with the model, we observed a plateau phase of memory in response to the combined input (Fig. 6B, “Experiment”). Moreover, in the absence of Pat1, the plateau phase was abolished and the memory dynamics largely resembled the sum of memory effects to the two single inputs (Fig. 6C, “Experiment”), consistent with the model prediction.

**Figure 6.**
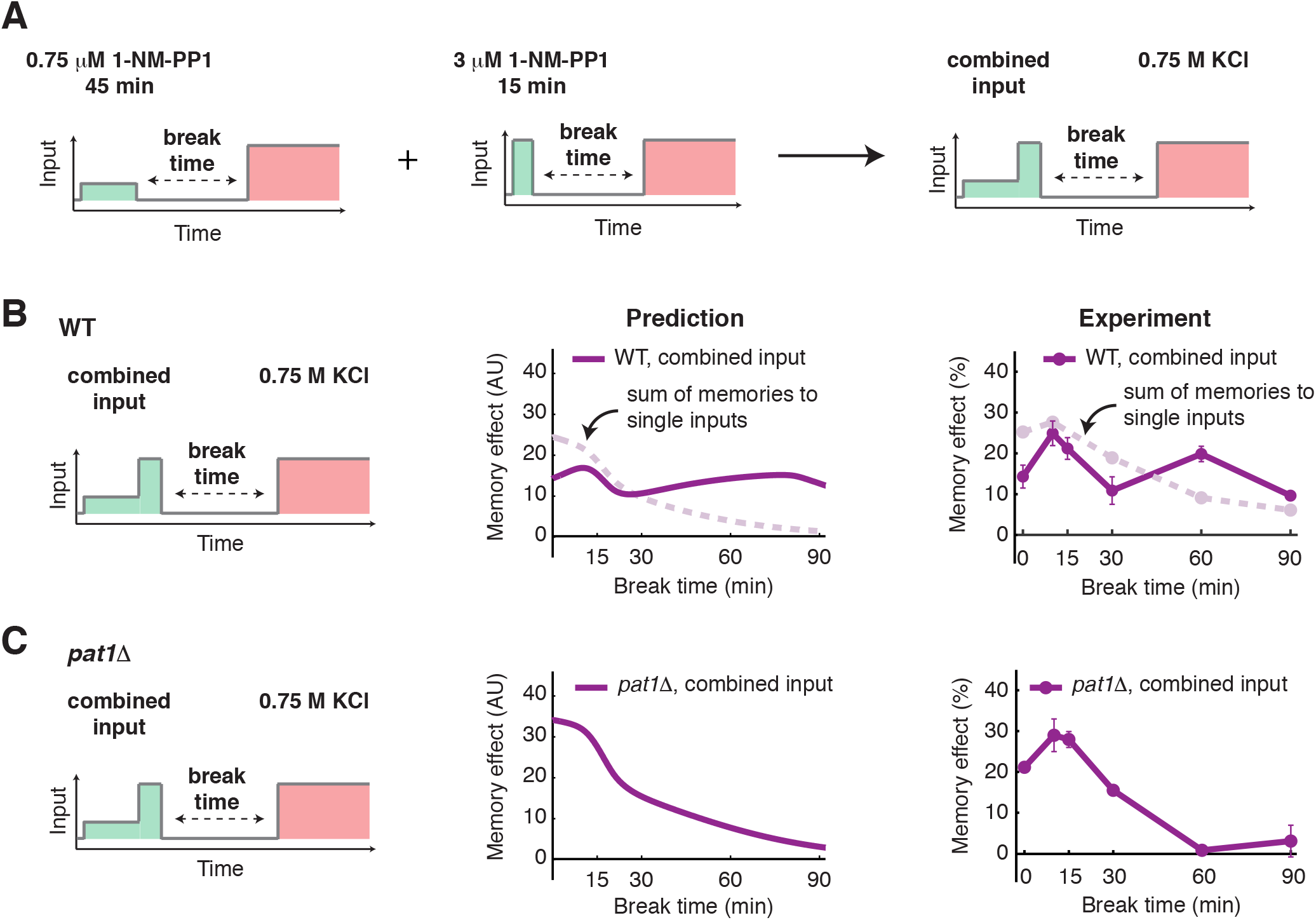
Model prediction and experimental validation. (A) A diagram illustrates the temporally combined input. Two single inputs (45 min, 0.75 *µ*M 1-NM-PP1 and 15 min, 3 *µ*M 1-NM-PP1) were applied sequentially to produce the combined input (45 min, 0.75 *µ*M 1-NM-PP1 followed by 15 min, 3 *µ*M 1-NM-PP1). (B) Model prediction and experimental validation of memory dynamics in WT in response to the combined input. Left panel, the schematic illustrates the treatment procedure. Middle panel, the plot shows the predicted memory dynamics in response to the combined input. Dashed line in light pink represents the sum of memories to two single inputs from Fig. 5A and 5B and is included in the plot for comparison. Right panel, the plot shows the experimentally measured memory dynamics in WT in response to the combined input. Dashed line in light pink represents the sum of memories to two single inputs from Fig. 2B and 2E and is included in the plot for comparison. (C) Model prediction and experimental validation of memory dynamics in *pat1*Δin response to the combined input.

These results validated our model and demonstrated its predictive power. Given that the memory-encoding processes are kinetically separated, the model-guided analysis open a powerful possibility to rationally design patterns of priming input and reprogram the temporal order of regulatory processes for generating desired forms of memory dynamics.

## Discussion

Cellular memory allows cells to adjust their responses to environmental cues based on their prior experience. In this study, we used GSR in yeast *Saccharomyces cerevisiae* as a model system and demonstrated that the memory effect on stress adaptation is biphasic, comprised of a fast decaying component (short-lived memory), mediated by post-translational regulation of the trehalose metabolism pathway, and a long-lasting component (long-lived memory), mediated by TFs and mRNP granules. These memory-encoding processes are mediated by PKA, a deeply conserved kinase with a central role in many molecular and cellular processes that is also associated with diverse diseases (Chiaradonna et al., 2008; Skalhegg and Tasken, 2000). Due to the functional pleiotropy of PKA, it has been difficult to target pharmacologically to achieve therapeutic specificity. Here we found that different PKA signaling dynamics, depending on the input amplitude and duration, could selectively induce specific downstream pathways or processes, leading to distinct memory dynamics. These results raise the possibility of perturbing the dynamics of signaling hubs for specific therapeutic outcomes (Behar et al., 2013; Li et al., 2017b).

Among the PKA-controlled processes, we want to emphasize the functional role of mRNP granules in maintaining cellular memory, which we revealed in this study. Cytoplasmic mRNP granules, such as PBs and SGs, play important roles in controlling the translation, degradation and storage of mRNAs upon environmental changes (Decker and Parker, 2012). Previous efforts have been focused on elucidating the biochemical and biophysical characteristics of mRNP granules (Mitchell and Parker, 2014). However, their functional roles in cell physiology remain largely unclear. Recent studies have begun to relate mRNP granules with cellular memory formation (Caudron and Barral, 2013; Chakrabortee et al., 2016; Dine et al., 2018; Standart and Weil, 2018). Our findings here provide independent strong evidence that the PKA-regulated mRNP granules contribute to a long-lasting memory of previous environmental challenges and facilitate the adaptation to future stresses. Further analyses will systematically determine the identities of these stored mRNAs that mediate the memory effect, the detailed mechanisms that direct these mRNAs to mRNP granules, and the prevalence of this mRNP-dependent memory effect in regulating other cellular responses, such as the hormetic effect on aging (Mattson, 2008).

In addition, through our modeling analysis, we obtained a quantitative understanding about the dynamics of cellular memory and the regulatory network that controls these dynamics, both of which remained largely missing from previous studies. The classic memory-generating networks feature positive feedback loops that give rise to bistability, underlying the mechanisms for irreversible processes such as cell-type differentiation (Chang et al., 2010; Wang et al., 2009; Xiong and Ferrell, 2003). In contrast, the network we identified is comprised of multiple parallel pathways with highly diversified inherent kinetics. This network architecture confers signal processing capability and plasticity in shaping memory dynamics, enabling cells to determine their memory patterns in response to rapidly changing environments. In particular, this system couples two low-pass filters with different thresholds to effectively separate the short-term responses to transient signals from the chronic ones to prolonged signals. Moreover, the system also features a PKA-regulated mRNP granule formation process, which represents a new network motif for biological information storage. The initiation of the storage depends on the input amplitude, whereas how long the storage can last depends on the amount of newly synthesized mRNAs being localized in the granules and hence depends on both the amplitude and the duration of priming inputs. In this way, mRNP granules enable cells to integrate the information about input amplitude and duration and tune the dynamics of their memory. Guided by the dynamic regulatory schemes revealed by modeling, we further designed priming input patterns to reprogram the temporal order of fast- and slow-acting processes in the network and reshape the memory dynamics. For future studies, advanced fluorescent reporters and imaging technologies (Aizer et al., 2014; Standart and Weil, 2018) could be employed to track the explicit spatiotemporal dynamics of key species in the model, such as mRNAs and mRNP granules, in single cells. These would enable us to constrain and improve our model and, ultimately, enhance the predictive power of the model. We anticipate that a quantitative and predictive understanding of memory control represents opportunities for broader and more effective use of priming treatments as a low-cost non-genetic approach for stress management in agriculture, biotechnology, and clinical intervention.

Finally, we want to highlight the biological relevance of our findings. We revealed that, because the molecular processes governing the two memory components have distinct kinetic properties, short- and long-lived memories could be selectively induced by different dynamics of priming inputs. Whereas a high amplitude transient input induces fast decaying memory that enables short-term stress resistance, a prolonged input elicits long-lasting memory conferring long-term stress resistance. This regulatory scheme is analogous to the fast responding innate immune response and the long-lasting adaptive immune response in mammals. Moreover, the mRNP granules are responsive to the amplitude and duration of inputs and can function as a knob to further tune the duration of the long-term memory component based on the input dynamics. Taken together, this integrated regulatory network enables cells to process the information of a previous stress encounter and determine how long they need to keep the memory about it. We speculate that this type of regulation may represent a strategy for cells to optimize resource allocation for future challenge preparation and may be widely applicable to organisms living in rapidly changing environments. Furthermore, given that this regulation is readily tunable, cells would be able to evolve their memory dynamics through natural selection to match the environmental fluctuations in their habitats.

## Acknowledgements

This work was supported by NIH R01 GM111458 (to N.H.) and NIH R35 GM128798 (to B.M.Z.).

## Author Contributions

Conceptualization, Y.J., Z.A., Y.L., B.Z., and N.H.; Methodology, Y.J., Z.A., Y.L., B.Z., and N.H.; Investigation, Y.J., Z.A., Y.L.; Formal Analysis, Y.J., Z.A., Y.L.; Writing - Original Draft, Y.J., Z.A., Y.L., and N.H.; Writing - Review & Editing, Y.J., Z.A., Y.L., B.Z., and N.H.; Resources, N.H.; Supervision, N.H.; Funding Acquisition, N.H.

## Materials and Methods

### Strain Construction

Standard methods for the growth, maintenance, and transformation of yeast and bacteria and for manipulation of DNA were used throughout. All *Saccharomyces cerevisiae* strains used in this study are derived from the W303 background (*MATa leu2-3,112 trp1-1 can1-100 ura3-1 ade2-1 his3-11,15 GAL*+ *psi*+ *ADE+*). The strains used in this study are listed in Table 1.

### Microfluidics

The previously reported Y-shape microfluidic device (AkhavanAghdam et al., 2016; Hao and O’Shea, 2012) with two inlets has been modified to accommodate three inlets on a single device and used in this study. The device fabrication and the setup of microfluidic experiments were performed as described previously (AkhavanAghdam et al., 2016; Hansen et al., 2015; Hao et al., 2013; Hao and O’Shea, 2012; Jiang et al., 2017).

### Time-lapse microscopy

All time-lapse microscopy experiments were performed using a Nikon Ti-E inverted fluorescence microscope with an Andor iXon X3 DU897 EMCCD camera and a Spectra X LED light source. A CFI Plan Apochromat Lambda DM 60X Oil Immersion Objective (NA 1.40 WD 0.13MM) was used for all experiments. Three positions were chosen for each microfluidics channel. For each position, phase contrast, YFP, mCherry, and iRFP images were taken sequentially every two minutes. When the acquisition of the image series started, cells loaded in the microfluidic device were maintained in synthetic complete medium (SC, 2% dextrose) for the first five minutes before the introduction of 1-NM-PP1. Media input was switched manually between SC medium, SC medium with 1-NM-PP1 and SC medium with KCl at the indicated time points. The exposure and intensity settings for each fluorescence channel were set the same as that used in our earlier work (AkhavanAghdam et al., 2016).

For the priming experiments, cells were inoculated from a YPD plate into 2 ml SC liquid media two days before the experiment. On the second day, saturated cells were diluted by 1:20,000 into fresh SC media and grown overnight to reach OD = 0.6. These exponentially growing cells were diluted by 1:2 and grown for another 2 hours before being loaded into the microfluidic devices.

### Image analysis

The images were processed using a custom MATLAB code for single-cell tracking and fluorescence quantification. The whole cell was segmented using the phase contrast images while the nucleus was segmented using the iRFP images. The cytoplasm was obtained by taking the difference of the whole cell and the nucleus. The nuclear to cytoplasmic ratios of Hog1-YFP were calculated using the mean fluorescence intensities of Hog1-YFP in the nucleus and in the cytoplasm. The ratios were subtracted by baseline which is the ratio right before KCl was introduced (close to 1) and then plotted against the time. The duration of Hog1 translocation for each condition was quantified using the full width at half maximum (FWHM) and used to calculate the memory effect for each break time, as illustrated in Fig. S1. We determine the sample size of our single-cell data based on similar studies published previously (AkhavanAghdam et al., 2016; Hansen and O’Shea, 2013; Hao et al., 2013; Hao and O’Shea, 2012).

### Trehalose assay

Trehalose level was measured using the trehalose assay kit (Megazyme). Using this assay, trehalose was converted into gluconate-6-phosphate, generating NADPH in a two-step reaction; the NADPH concentration can be determined by measuring the absorbance at 340 nm. To determine trehalose concentrations, 13 mL of yeast culture at OD≈5 was harvested and put on ice for 5 minutes before centrifuged for 5 min at 4 °C. Cells were then washed with 0.1 M phosphate buffer (pH 5.9) to remove glucose in media. After the wash, cells were resuspended in 1 ml 0.25 M Na_2_CO_3_ solution and OD was measured. Additional Na_2_CO_3_ solution was added to make the cell densities (OD) the same for 0 and 20 min samples. Samples (∼1 mL) were boiled for 20 min to release intracellular trehalose. After cooling, the samples were centrifuged at 12 000 g for 3 min to remove cell debris. Two aliquots (300 µl) of supernatant were transferred to two new tubes with one tube for total glucose level and the other one for pre-existing glucose level. The following regents were then added to the cell lysates sequentially: 150 µl acetic acid (1 M), 650 µl distilled water, 100 µL imidazole buffer (2 M imidazole, 100 mM magnesium chloride and 0.02% w/v sodium azide; pH 7.0), 50 µL NADP 1 /ATP (12.5 mg/mL NADP+ and 36.7 mg/mL ATP) and 10 µL suspension of HK/G-6-PDH (425 U/mL hexokinase and 212 U/mL glucose-6-phosphate dehydrogenase), 10 µL trehalase (490 U/mL). The mixtures were incubated at room temperature for 5 min for the reactions. Absorbance at 340 nm was recorded to determine the trehalose concentration in solution first. To estimate the intracellular concentration, we assumed that cell density at OD=1 is 1×107 cells/ml and yeast cell volume is 42 fl. Pre-existing glucose was determined in a control reaction without added trehalase.

### Live-cell mRNA visualization

The MS2-CP strains for mRNA visualization were constructed as described previously (Zid and O’Shea, 2014). The promoter and the coding region of genes of interest (e.g. *DCS2*) were amplified by PCR and then inserted into a template vector, which contains 12× MS2 loop sequences in the integration vector pRS305. The plasmid was linearized by EcoRV and integrated into W303 background yeast strain with PKA analog-sensitive mutations (NH084) at the *LEU2* locus. A plasmid that constitutively expresses MS2 coat proteins fused with GFP driven by *MYO2* promoter (Zid and O’Shea, 2014) was also integrated into the same strain at *HIS3* locus. To visualize the colocalization of mRNAs and PBs, a pRS304 plasmid that expresses *DCP2*-mCherry under the native *DCP2* promoter was integrated into the same strain at *TRP1* locus (NH0857).

To perform live-cell mRNA visualization, cells were cultured to OD 0.6 and then loaded into microfluidic devices for time-lapse microscopy. For each position, phase contrast, mCherry, GFP and iRFP images were taken sequentially every two minutes. When the image acquisition started, cells were maintained in SD media for the first five minutes to obtain a baseline for each fluorescence channel prior to the introduction of 3 μM 1-NM-PP1 treatment.

The exposure and intensity settings for each channel were set as follows: GFP 200 ms at 9% lamp intensity, mCherry 1s at 10% lamp intensity, and iRFP 300 ms at 15% lamp intensity.

### Computational Modeling

Our model focuses on the PKA-dependent memory-encoding network, comprised of the experimentally-identified processes that regulate the levels of metabolites or proteins needed for enhancing stress adaptation, including trehalose and stress resistance gene products. The input of the model is the PKA inhibition signal (priming input). For the model output, we assume that the memory effect linearly depends on the sum of the amounts of trehalose and stress resistance gene products in most of the kinetic regimes, unless the substance concentrations reach extremely high levels. This assumption leaves out the explicit inclusion of the downstream Hog1 pathway in our model and simplifies our analysis. The network consists of three molecular processes, trehalose metabolism, stress resistance gene transcription and mRNP granule formation, all of which are PKA regulated.

For trehalose metabolism, PKA regulates both trehalose production and degradation by phosphorylating Tps1 and Nth1, respectively. More specifically, PKA-mediated phosphorylation inhibits Tps1 activity and enhances Nth1 activity based on the previous reports. Because the regulation is primarily through phosphorylation, we assume that it is a fast process. For stress resistance gene expression, PKA regulates mRNA transcription through transcription factors and regulates mRNA translation and degradation through mRNP granules. More specifically, the PKA inhibition input activates transcription factors and mRNP granules. Once the mRNP granules are activated, a portion of the newly synthesized mRNAs is stored in mRNP granules where their translation and degradation are paused. Upon input removal, these mRNAs are slowly released, resuming their translation and degradation. Because gene transcription is a multi-step process, we assume that it is a relatively slow process; in contrast, because PKA regulates mRNP granules through phosphorylation, we assume that it is a relatively fast process. It should be noted that the level of Tps1 is also dependent on PKA-regulated gene expression, resulting in a connection between the trehalose metabolism pathway and the gene expression process. Based on the experimental observations, these three processes have different dependence on input amplitude and duration. Trehalose metabolism and mRNP granules can only be activated in response to high-amplitude inputs; by contrast, mRNA transcription can be induced by low-amplitude input but a prolonged duration is needed.

Computational modeling and all the simulations were done using the MATLAB. The model contains 13 variables and 27 independent parameters. The function “lsqnonlin” was used for data fitting. The data of three dynamic inputs and three mutants (*tps1Δ, pat1Δ, msn2/4*Δ*yap1Δ*) were used for data fitting (Fig. S5). To highlight the role of mRNP granules, time points for long-lived memory plateau (Fig. 1F) were weighted by 20-fold for data fitting. Fitting starting with completely random guesses failed. Therefore, we first manually chose parameter values that can qualitatively capture the data and then used these manually-chosen parameters as the initial guesses for computational fitting. In order to overcome the local minimum, we also tested 10 random sets of initial guesses which are randomly chosen within 8 fold (1/8 to 8) of our first set of guesses and compared the squared norm of the residuals of the final fitting results and then selected the best-fit set of parameter values.

The initial conditions are provided in Table S1. Reactions and rate constants are provided in Table S2.

### Model Equations

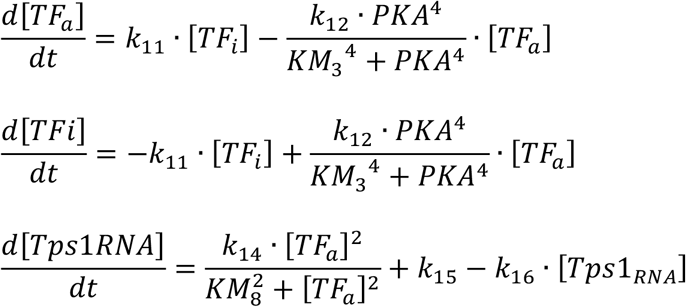

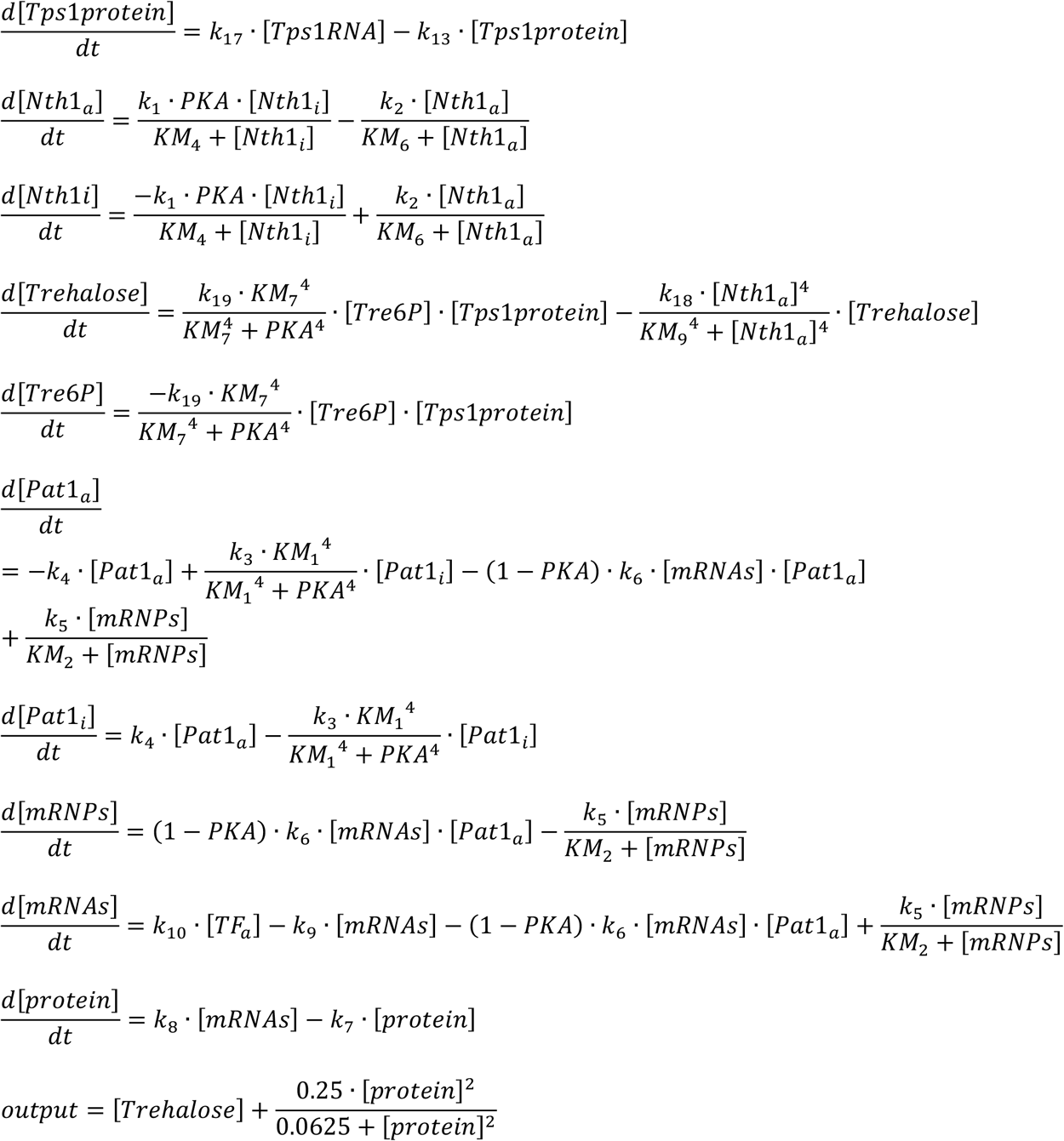

### Strains used in this study

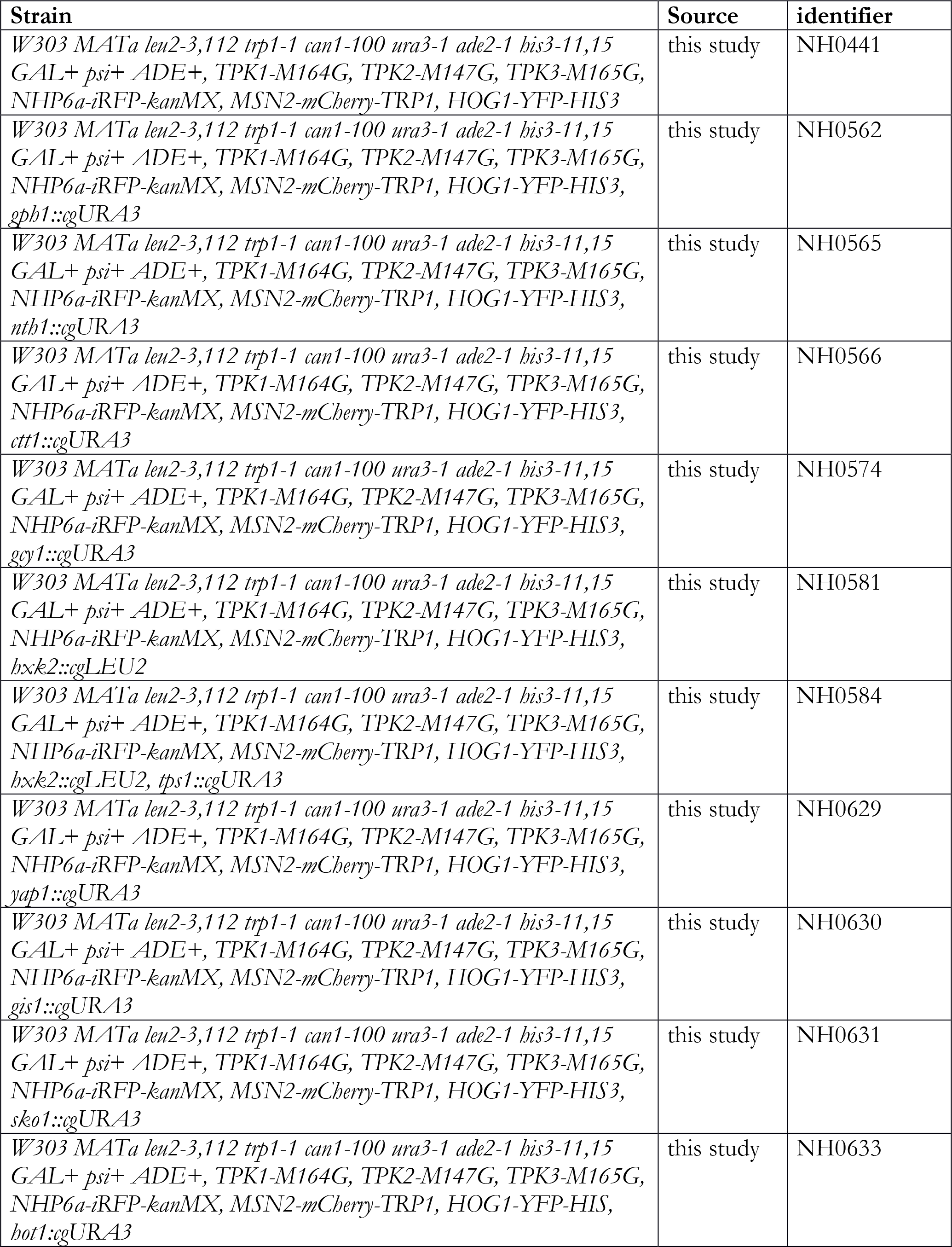

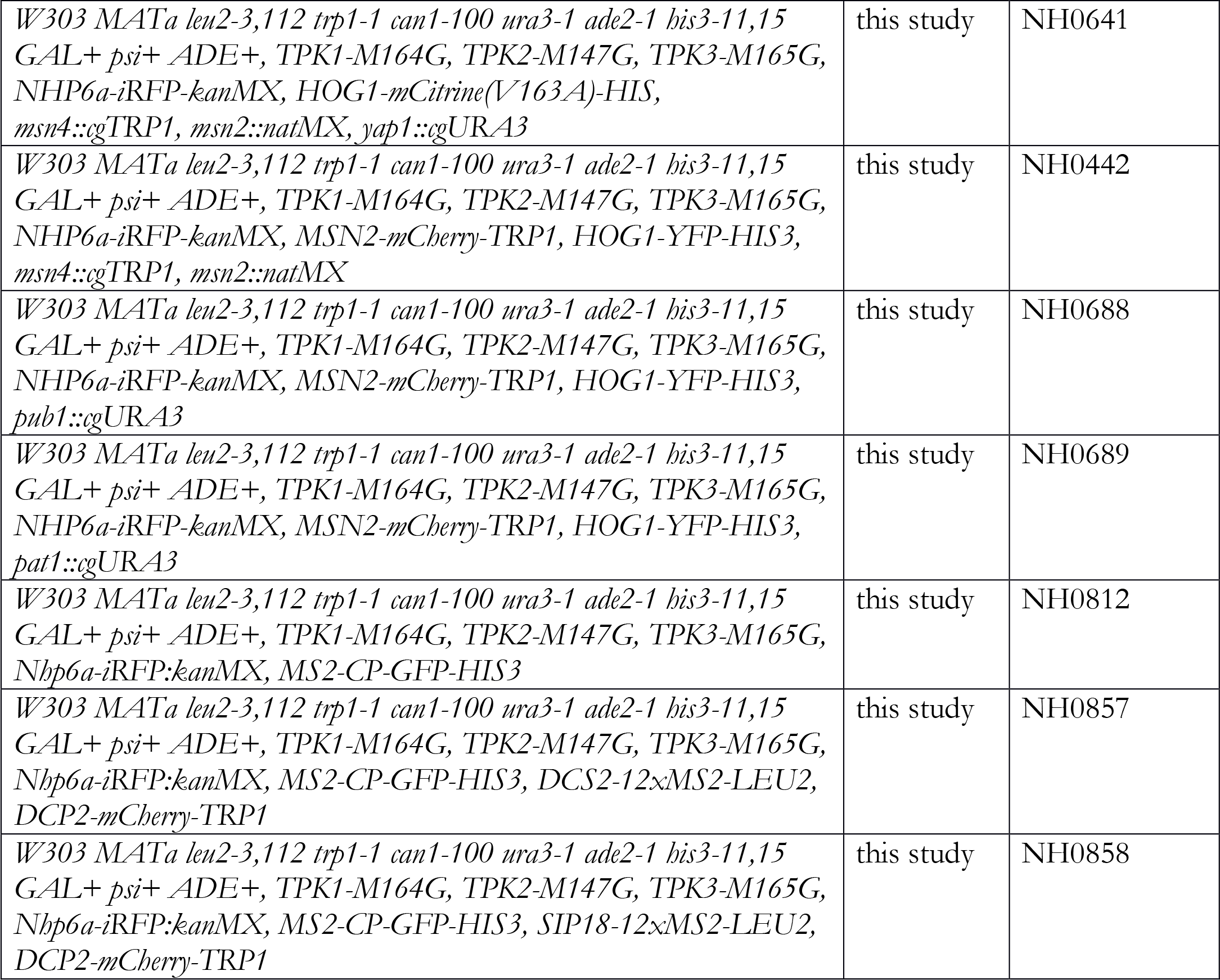

**Table S1.**
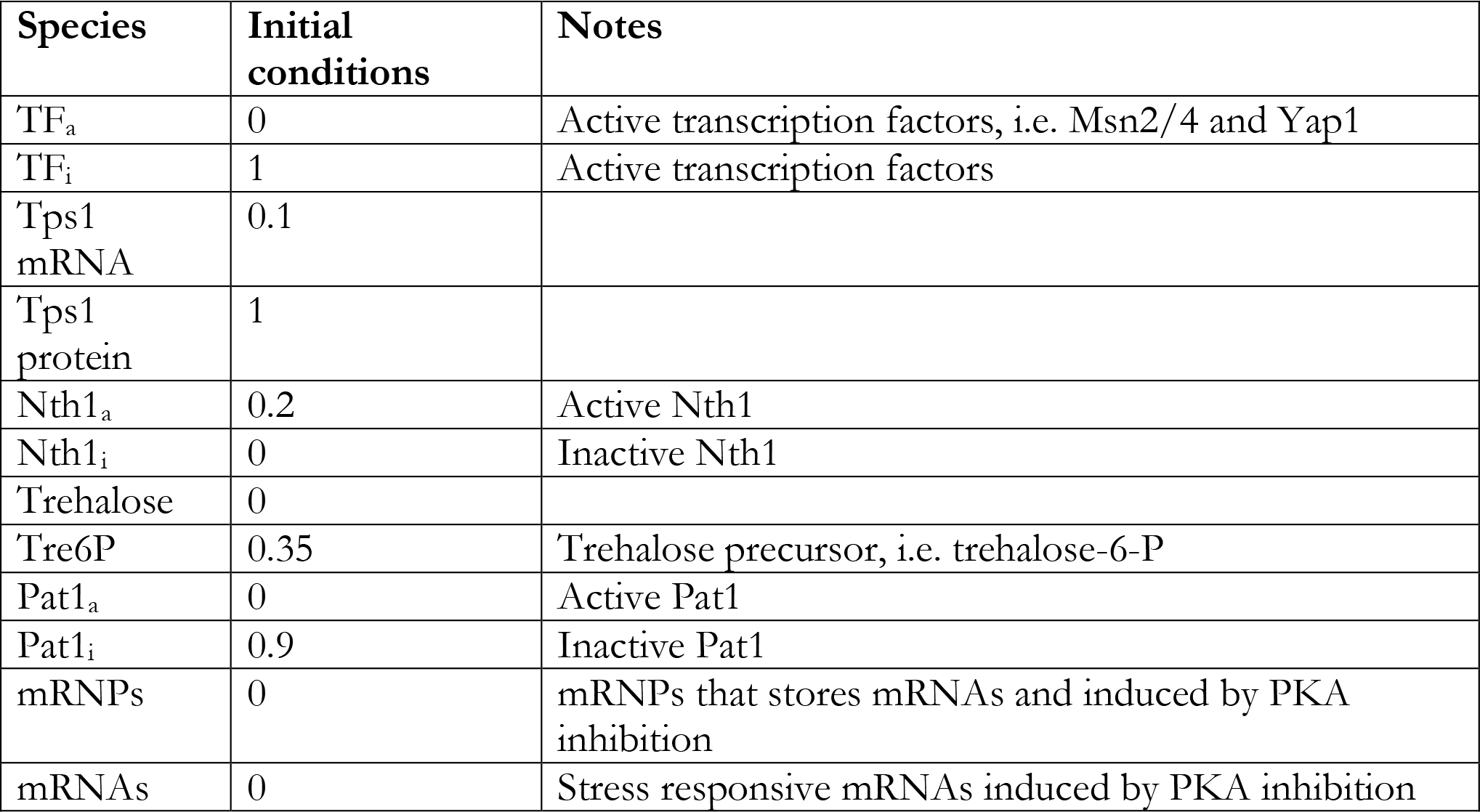
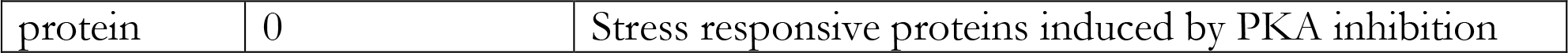
Initial conditions of all species in the model.

**Table S2.** Best-fit parameter values used in the model.

| Names | Parameters | Values |
| --- | --- | --- |
| PKA activity | PKA | no inhibitor: 1<br>3μM1-NM-PP1: 0<br>0.75μM1-NM-PP1: 0.25 |
| Transcription factor activation rate | k<sub>11</sub> | 0.009 min<sup>-1</sup> |
| Transcription factor deactivation rate | k<sub>12</sub> | 50 min<sup>-1</sup> |
| Transcription factor deactivation equilibrium constant | KM<sub>3</sub> | 1.3 |
| Basal level of Tps1 mRNA transcription | k<sub>15</sub> | 0.01 min<sup>-1</sup> |
| Tps1 mRNA transcription rate | k<sub>14</sub> | 0.4 min<sup>-1</sup> |
| Tps1 mRNA degradation rate | k<sub>16</sub> | 0.05 min<sup>-1</sup> |
| Tps1 translation rate | k<sub>17</sub> | 0.005 min<sup>-1</sup> |
| Tps1 degradation rate | k<sub>13</sub> | 0.005 min<sup>-1</sup> |
| Tps1 activation equilibrium constant | KM<sub>6</sub> | 0.01 |
| Tps1 mRNA transcription equilibrium constant | KM<sub>7</sub> | 0.04 |
| Nth1 activation rate | k<sub>1</sub> | 0.51 min<sup>-1</sup> |
| Nth1 deactivation rate | k<sub>2</sub> | 0.5 min<sup>-1</sup> |
| Nth1 activation equilibrium constant | KM<sub>4</sub> | 0.00001 |
| Nth1 deactivation equilibrium constant | KM<sub>5</sub> | 0.00001 |
| Trehalose production rate | k<sub>19</sub> | 0.02 min<sup>-1</sup> |
| Trehalose decay rate (rate converting to downstream products) | k<sub>18</sub> | 0.41 min<sup>-1</sup> |
| Trehalose decay equilibrium constant | KM<sub>8</sub> | 0.2 |
| Pat1 activation rate | k<sub>3</sub> | 5 min<sup>-1</sup> |
| Pat1 deactivation rate | k<sub>4</sub> | 0.5 min<sup>-1</sup> |
| Pat1 activation equilibrium constant | KM<sub>1</sub> | 0.01 |
| mRNPs formation rate | k<sub>6</sub> | 5 min<sup>-1</sup> |
| mRNPs disassembly rate | k<sub>5</sub> | 0.01 min<sup>-1</sup> |
| mRNPs disassembly equilibrium constant | KM<sub>2</sub> | 0.01 |
| mRNA transcription rate | k<sub>10</sub> | 0.16 min<sup>-1</sup> |
| mRNA degradation rate | k<sub>9</sub> | 0.02 min<sup>-1</sup> |
| Stress resistance protein translation rate | k<sub>8</sub> | 3.2 min<sup>-1</sup> |
| Stress resistance protein degradation rate | k<sub>7</sub> | 4 min<sup>-1</sup> |

