## Supplementary Figures for "A Protein Kinase A-Regulated Network Encodes Short- and Long-Lived Cellular Memory"

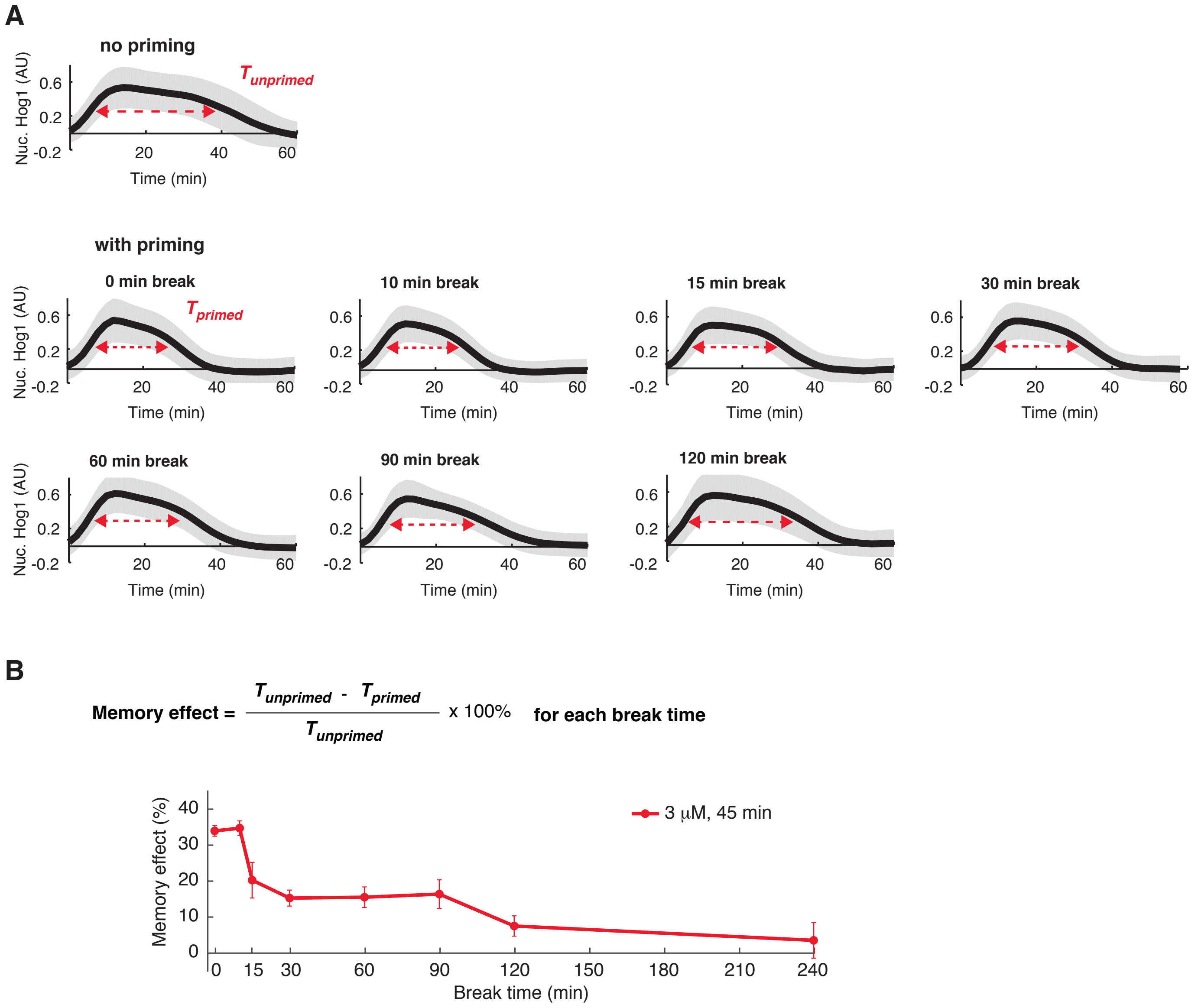


**Figure S1. Measurement and Quantification of memory effects as a function of the break time.** (A) Quantified time traces of Hog1 nuclear translocation with different break times. The x-axis indicates the time that sustained 0.75 M KCl treatment has been introduced. The y-axis is the ratio of the mean fluorescence intensities in the nucleus and in the cytoplasm. The gray shaded areas represent the standard deviations of single-cell traces, while the black curves represent the means. Baseline has been subtracted in all traces. The duration (adaptation time, T) is defined as the full width at half maximum (FWHM, red dashed line with arrows) for each time trace, which is unaffected by any normalization in the y-axis. The first of the eight subpanels shows the results of the experiments without priming. The rest seven subpanels are the results of the experiments with 45 min, 3 μM 1-NM-PP1 and different break times before the addition of 0.75 M KCl. (B) Quantification of memory effects based on adaption times. "Memory effect" is defined with the indicated equation and can be calculated based on adaptation times obtained from the time traces in Panel A. The resulting "memory effects" are plotted vs the break times. It should noted that the memory effects goes to 0 when the priming treatment has no effect and the Hog1 translocation is not affected when compared to the unprimed experiments, while it goes to 100% when the there is no Hog1 translocation in response to KCl, which indicates the cells are perfectly prepared for the KCl stress.

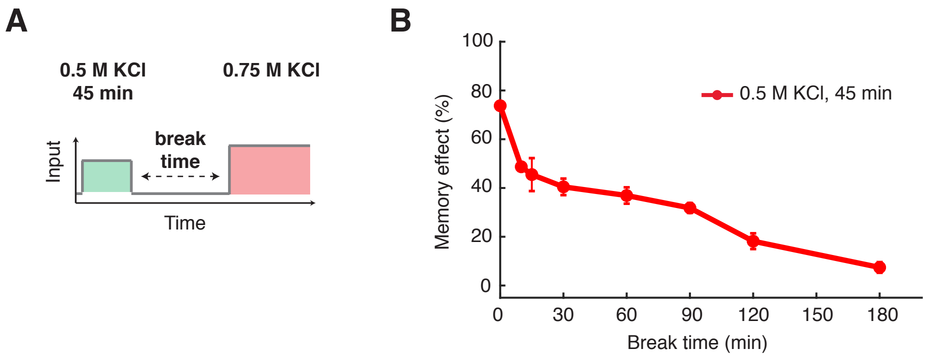


**Fig. S2. Memory dynamics in response to 45 min 0.5 M KCl priming input.** (A) The schematic illustrating the treatment procedure of the priming experiment - 45 min, 0.5 M KCl priming input. (B) The plot shows the relationship of memory effect versus break time. Error bars – standard error of the mean (SEM).

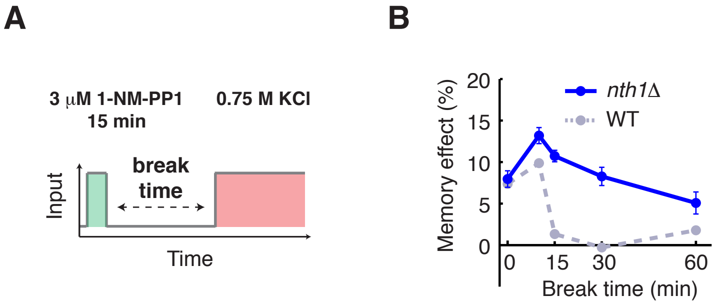


**Figure S3 . Short-lived memory is regulated by the trehalose degradation enzyme, Nth1.** (A) The schematic illustrating the treatment procedure of the priming experiment - the high-amplitude transient priming input (15 min, 3 *μ*M 1-NM-PP1). (B) Memory dyanmics in *nth1Δ* in response to the high-amplitude transient priming input . Dashed lines in grey blue represent the memory dynamics in WT (from Fig. 2B) and are included in the plots for comparison.

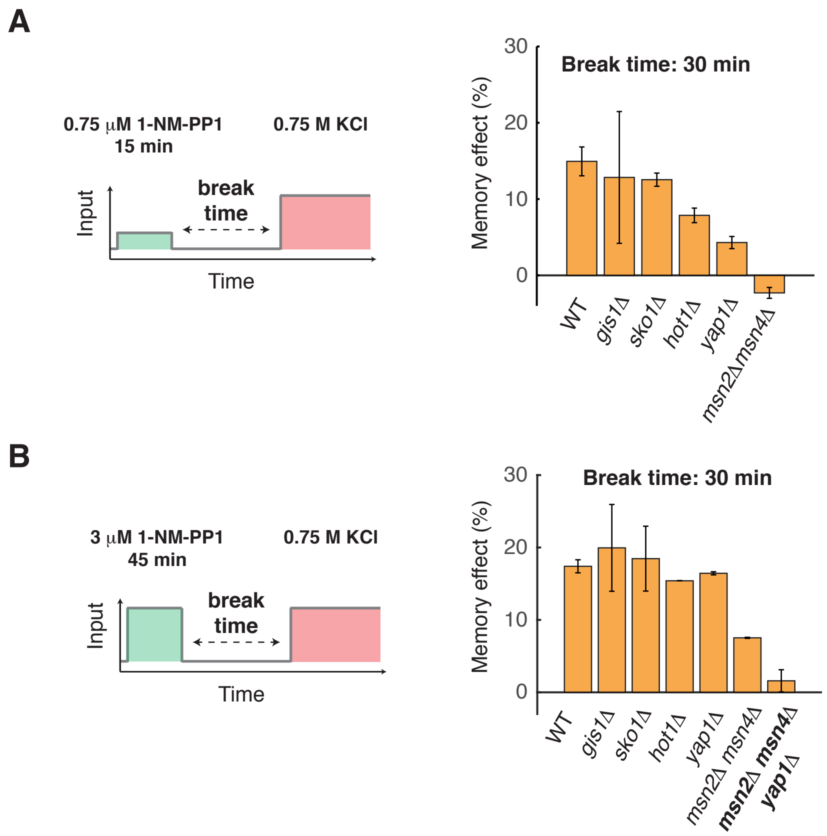


**Figure S4. Identification of TFs Msn2/4 and Yap1 in mediating long-lived memory.** (A) Bar graph shows the memory effects in response to (A) the low-amplitude prolonged priming input (45 min, 0.75 $\mu$M 1-NM-PP1) or (B) the high-amplitude prolonged priming input (45 min, 3 $\mu$M 1-NM-PP1), with a 30 min break time in WT and mutant strains.

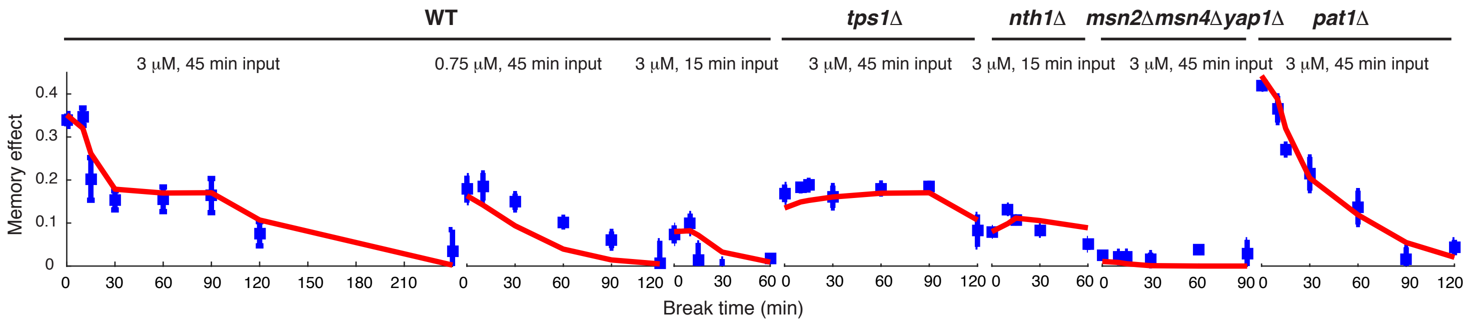


**Figure S5. Model fitting with all the experimental data.** The blue squares show the experimental data with the error bars representing SEMs. The red lines are the modeling results with the best-fit parameter values. Seven data sets were used for data fitting. The first three data sets are from Figs.1 and 2, which show the memeory decay with three different inputs. The *tps1*Δ, *msn2/4*Δ *yap1*Δ, and *pat1*Δ data are from Figs. 3 and 4. The *nth1*Δ data are from Fig. S3. The fitting method is described in the Methods.

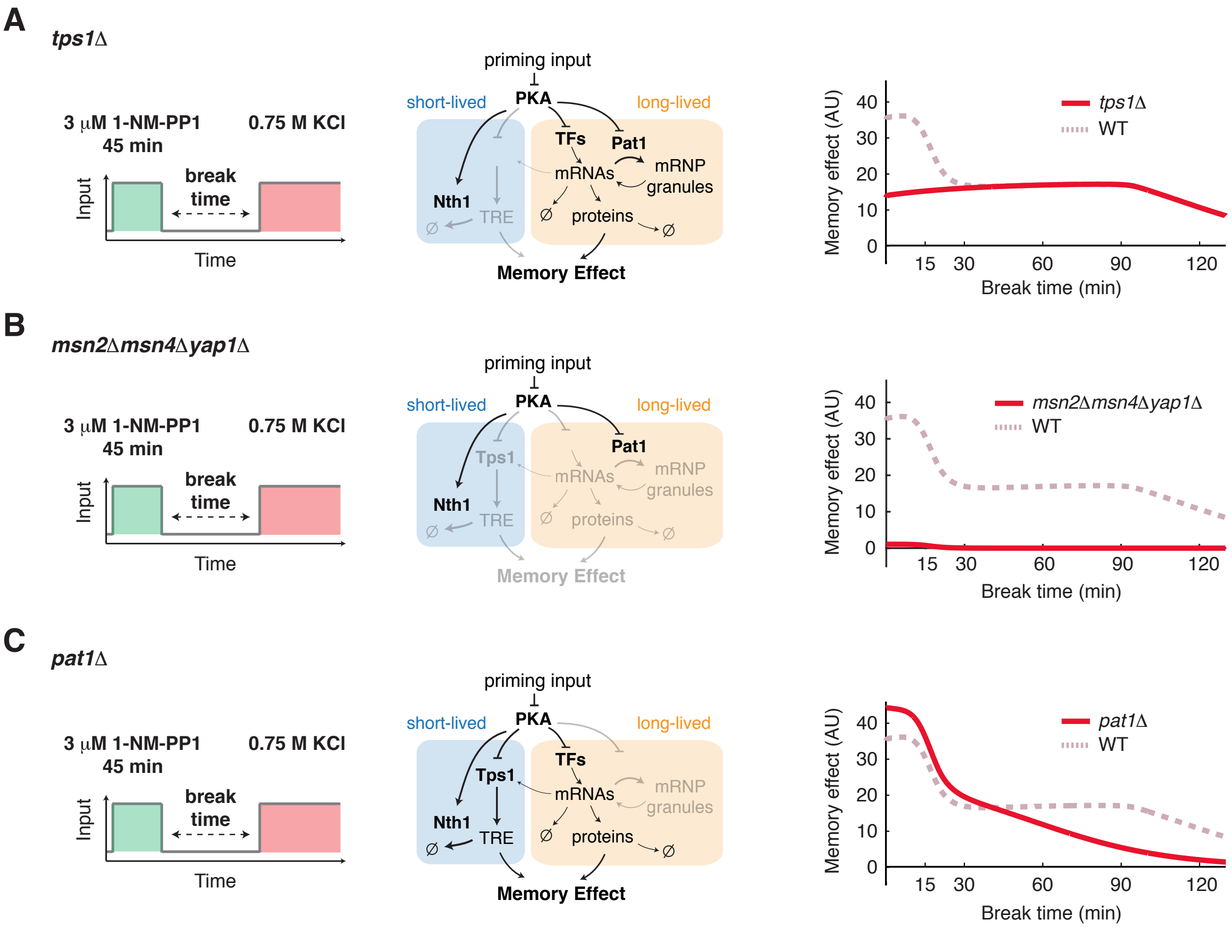


**Figure S6. Computational modeling reproduces the modulation of memory dynamics in the absence of key regulatory factors.** (A) Model simulation of memory dynamics in the absence of Tps1. Left panel, the schematic illustrates the treatment procedure (45 min, 3 $\mu$M 1-NM-PP1). Middle panel, the diagram highlights the part of the network that remains activated in the absence of Tps1 (black – activated; light gray – not activated). Right panel, the plot shows the simulated dynamics of memory effect. Dashed line in dark pink represents the simulated dynamics of WT from Fig. 5C and is included in the plot for comparison. (B) Model simulation of memory dynamics in the absence of Msn2/4 and Yap1. (C) Model simulation of memory dynamics in the absence of Pat1.
